# Game of full siblings in Mendelian populations

**DOI:** 10.1101/2022.12.13.520274

**Authors:** József Garay, András Szilágyi, Tamás Varga, Villő Csiszár, Tamás F. Móri

## Abstract

We adapt the concept of evolutionary stability to familial selection when a game theoretic conflicts between siblings determines the survival rate of each sibling in monogamous, exogamous families in a diploid, panmictic population. Similarly to the classical evolutionary game theory, the static condition of evolutionary stability of mixed Nash equilibrium implies the local stability of the genotype dynamics, in spite of that the mating table based genotype dynamics is not a replicator dynamics.

We apply our general result to the case where a matrix game determines the survival rate of siblings. In our numerical studies we consider the prisoner’s dilemma between siblings, when the cooperator and defector behaviour are unequally determined by a recessive-dominant allele pair at an autosomal locus. When the prisoner’s dilemma game is strict (cf. iterated one) and the cooperator phenotype is recessive resp. dominant, then the cooperator and defector phenotypes are the unique stable phenotypes, respectively. When the prisoner’s dilemma game is not strict, both phenotypes coexist, independently of the genotype-phenotype mapping. However, the frequencies of the phenotypes are different according to which phenotype is dominant.

## 1. Introduction

We start out from Haldane’s (1924) “*familial selection*” situation, where the size of the family is strictly limited by the food and more newborns are produced than can survive to enter the struggle among the members of the same family. Here we consider a slightly different selection situation, as we do not assume that each family in a population contributes the same number of adults in the next generation (e.g. Theodorou and Couvet, 2003), and every sibling’s survival rates depends on their siblings’ behaviour. In other word, here we consider frequency dependent, survival game-theoretic conflicts between siblings (cf. Garay 2009; Garay and Varga, 2011) within monogamous and exogamous families (Garay et al., 2019, 2023). We emphasize that the game theoretical conflict is well mixed between full siblings, but is not in the whole population.

During sexual reproduction haploid and diploid life cycles follow each other. Thus there are two ways to model the evolution in diploid sexual populations: follow either the haploid stage (gamete types) or the diploid stage (genotypes) from generation to generation.

### Gene centred models

These kinds of models are usually based on gene pool approach. The diploid population is at genetic equilibrium and the variable of the models is the allele distributions (e.g. Fisher, 1958; Maynard Smith, 1989; Nagylaki, 1994).

### Genotype centred models

When the fertilization is internal and the juveniles’ survival rate depends on the mating pair, then the genotype centred population genetic model is more reasonable. There the state space of the model is the set of diploid genotype distributions (e.g. Cavalli-Sforza and Feldman, 1978; Garay et al., 2019; Haldane and Jayakar, 1965; Hull, 1964; King, 1965; Maynard Smith, 1980; Scudo and Ghiselin, 1975; Uyenoyama and Feldman, 1980; Uyenoyama et al., 1981). For instance, in the simplest case (one autosomal locus with only two alleles), although under panmixia the diploid embryos are exactly in Hardy–Weinberg proportions, but the family dependent survival selection can radically change this proportions in adults. Thus the Hardy–Weinberg equilibrium proves to be useful neither in model formulation nor in the analysis of the models (see e.g. Nagylaki, 1994). We mention two additional advantages of the genotype centred population models. Firstly, the classical Darwinian fitness is defined as the growth rate of the diploid phenotype. Consequently, the genotype centred model is the nearest to the classical Darwinian view. Secondly, kin selection theory focuses on the diploid individuals, the cost and benefit from their interactions, and the genetic relatedness between them (Hamilton, 1964). In this sense, Hamilton’s rule and the notion of inclusive fitness is also genotype centred.

The genotype centred population genetics models have already been used to study the evolution of altruism within the families (e.g. Cavalli-Sforza and Feldman, 1978; Garay et al., 2019, 2023; Maynard Smith, 1980; Uyenoyama and Feldman 1980; Uyenoyama et al., 1981). We recall some results about these population genetics models, which can be considered as preliminaries of the present paper. Firstly, the fixation probability of the altruistic behaviour depends on the genotype-phenotype mapping (Garay et al., 2019; Toro et al., 1982; Uyenoyama and Feldman, 1980). Secondly, in the presence of additive cost and benefit functions the population genetics models give the same results that Hamilton’s method gives (Garay et al., 2019, 2023; Hamilton, 1964), but does not in the non-additive situation (Cavalli-Sforza and Feldman, 1978; Garay et al., 2019, 2023). All of these published models focus on the evolutionary stability of the monomorphic altruistic population. Contrary to that, here we concentrate on the evolutionary stability of a polymorphic population, i.e. we are looking for the coexistence of two different phenotypes. This is the main novelty of the present paper. Thus, we focus on the following two problems: 1. What are the conditions of the existence of a stable mixed genotype distribution? 2. Do these conditions imply the dynamical stability of the corresponding genotype dynamics?

Furthermore, the altruism can be considered as a one-person game, where an actor has two pure strategies (altruistic or selfish) and the recipient has no strategy. Then the following question arises: does Hamilton’s rule remain valid, or does not if the recipient also has strategy, i.e. the interaction between siblings can be given by a two-person game? We consider two dominant-recessive alleles at an autosomal locus, which unequally determine the phenotypes, namely two pure strategies, and we present numerical examples when a matrix game describes the interaction between the siblings. Since one of the most popular and widely studied game theoretical conflicts is the prisoner’s dilemma (Poundstone, 1993), we will also concentrate on this dilemma.

## 2. General mating table based model

### 2.1. Familial selection

Here we start out our earlier introduced model (Garay et al., 2019). We assume that the phenotypes are uniquely determined by genotypes. In monogamous families the parents’ genotypes determine the genotype distribution of their offspring, thus the phenotypes of the offspring, too. Between siblings, there is a game theoretical conflict. The main point here is that the interactions take place within the family only. Consequently, the survival rate of each juvenile does not depend on the phenotype distribution of the whole population.

Now we give the details of the biological system we will use (Garay et al., 2019). We consider a diploid population of (large) size *N* with sex ratio half-half, with no sexual selection, i.e. the females and males differ only in their sex. In other words, we consider autosomal loci here. The species is monogamous with internal fertilization. We consider *panmixia,* i.e. well-mixed reproductive pair formation between genotypes in the whole population. Thus the probability of two genotypes’ mating depends only on their frequencies in the population, so there is no assortative mating. The species is monogamous, so *N*/2 couples are formed at random, and each couple breeds a fixed number *n* of offspring. We emphasize that if the population is large enough, then the random mating excludes inbreeding. Thus, the degree of relatedness between full siblings is ½. Since our model allows that the game is a survival game, the diploid parental population is not in Hardy–Weinberg equilibrium. The generations are not overlapped.

For the convenience of the reader, we start the simplest genetic situation, namely, we consider two alleles, but we allow arbitrary interactions between siblings. In this special case we have the mating table given in Table 1. We note that applying our general model we also consider the two allele case. Let *N* be the population size of the whole parent population, then 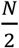 is the number of parent pairs. Let *x* = (*x*_1_, *x*_2_, *x*_3_) be the frequency vector of genotypes *G*_1_, *G*_2_ and *G*_3_ in the present generation. Then

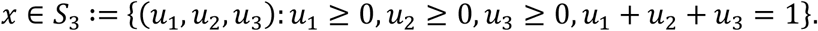

**Table 1.** Genotype survival table based on the mating table

| Genotypes of parents | Average number of couples | Probability distribution of offspring, and Average number of survived family members, for genotypes $G_1, G_2, G_3$ | | |
| --- | --- | --- | --- | --- |
| | | $G_1 = ([a],[a])$ | $G_2 = ([a],[A])$ | $G_3 = ([A],[A])$ |
| $G_1 \times G_1$ | $\frac{N}{2} x_1^2$ | 1<br>$n_{1(11)}$ | 0<br>0 | 0<br>0 |
| $G_1 \times G_2$ | $N x_1 x_2$ | $\frac{1}{2}$<br>$n_{1(12)}$ | $\frac{1}{2}$<br>$n_{2(12)}$ | 0<br>0 |
| $G_1 \times G_3$ | $N x_1 x_3$ | 0 | 1<br>$n_{2(13)}$ | 0 |
| $G_2 \times G_2$ | $\frac{N}{2} x_2^2$ | $\frac{1}{4}$<br>$n_{1(22)}$ | $\frac{1}{2}$<br>$n_{2(22)}$ | $\frac{1}{4}$<br>$n_{3(22)}$ |
| $G_2 \times G_3$ | $N x_2 x_3$ | 0<br>0 | $\frac{1}{2}$<br>$n_{2(23)}$ | $\frac{1}{2}$<br>$n_{3(23)}$ |
| $G_3 \times G_3$ | $\frac{N}{2} x_3^2$ | 0<br>0 | 0<br>0 | 1<br>$n_{3(33)}$ |

Since the parental population is not at Hardy–Weinberg equilibrium, the relative frequency of parental genotype *G_i_* does not follow from the relative frequencies of the alleles of the genotype in question. According to the random mating, the relative frequency of families with parents of genotypes *G_i_* and *G_j_* is given by 2*x_i_x_j_*.

Let *n*_*k*(*ij*)_ denote the number of surviving offspring of genotype *G_k_* in a monogamous family with parents of genotypes *G_i_* and *G_j_*. We note that *n*_*k*(*ij*)_ can be determined either by pairwise interactions or by synergetic group effect (Garay et al., 2023), or in any other way.

For instance, consider a family with parents with of genotypes *G*_1_ and *G*_2_. According to the random segregation of alleles (or chromosomes), this family has *G*_1_ type offspring with probability ½, and *G*_2_ type offspring with probability ½, as well. Thus the genotype distribution in this family is a binomial one, i.e. 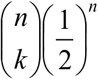 is the probability that in this family there are exactly *k* offspring of genotype *G*_1_

#### Remark.

Observe that the basic assumption of replicator dynamics (i.e. each genotype is produced by the same genotype only) is not valid, due to Mendelian inheritance. For instance, a homozygote offspring can be produced by heterozygote parents, too.

Based on Table 1, the total numbers of individuals of genotypes *G*_1_, *G*_2_ and *G*_3_ in the next generation are the following:

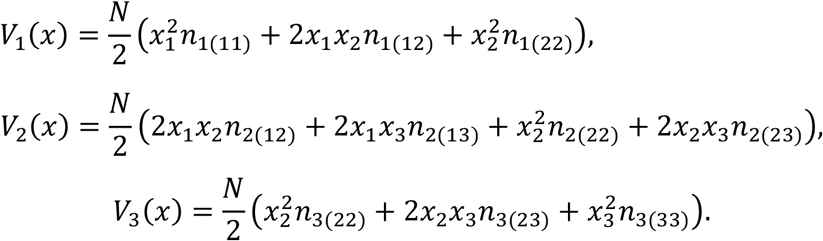

Hence, if *x* ∈ int *S*_3_ (thus *x_i_*, > 0 for *i* ∈ {1, 2, 3}) the average growth rate (fitness) of genotype *G_i_* is

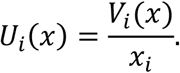

The average growth rate of the population at *x* ∈ int *S*_3_ is

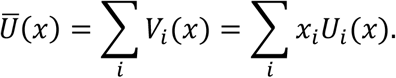

We emphasize that, though at first glance this genotype centred model seems to be a phenotypic one, the production rate *n*_*k*_(*ij*) of offspring with genotype *Gk* in a monogamous family with parents of genotypes *G_i_* and *G_j_* clearly depends on the genetic system. After considering the case of one autosomal locus with two alleles (see Table 1), we can easily treat more complex situations with an arbitrary number *m* of genotypes, see Appendix A.

### 2.2. Evolutionary stability of genotype distributions

Now we ask when a genotype distribution is an *evolutionarily stable genotype distribution* (ESGD) in an arbitrarily large sexual population. Following Maynard Smith and Price (1973), we say that a homozygote is ESGD if the rare mutant gene cannot invade the resident population (Garay at al., 2019, 2023). Mathematically, this means that the (relative) frequency of the resident homozygote increases from generation to generation, provided mutation is sufficiently rare. However, the original idea of Maynard Smith and Price (1973) has no direct meaning when there is polymorphism in the genotype (with pure phenotype) population. Based on the dynamical view of the stability we can say that a polymorphic stage is an ESGD if, when the frequency of a genotype increases by mutation (or other type of perturbation), then the system goes back to the ESGD. A static formulation this idea is the following. A genotype distribution is called an ESGD if the mean production rate (fitness) of the whole genotype population (see right-hand side of inequality (1) below) is less than that of the genotype subpopulation at ESGD (in the left-hand side of inequality (1)). Note that here we use the idea of our condition for the evolutionary stability of allele distributions (Garay and Varga, 2003).

#### Definition 1.

*A genotype distribution x*\* ∈ int *S_3_ is evolutionary stable if*

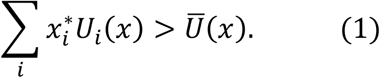

*provided that x is sufficiently close to x*.*

In another form,

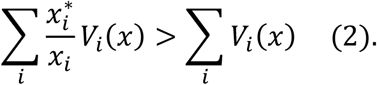

Note that *V_i_* (*x*)/*x_i_* gives the number of genotype *G_i_* offspring that a single individual of the same genotype in the parental generation has. Thus *V_i_* (*x*)/*x_i_* can be considered as the fitness of genotype *G_i_*. As defined above, *x*\* ∈ int *S*_3_ is an ESGD if and only if

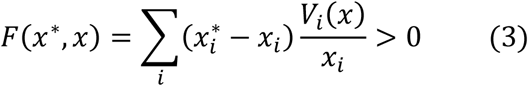

whenever *x* is close enough to *x*.*

#### Remark.

Definition 1 has two interpretations. In the monomorphic setup each individual can use mixed strategies (thus the evolutionarily stable phenotype chooses from the strategies randomly, with probabilities given by the ESGD). In the polymorphic setup each individual can use no more than one strategy, thus only a mixed population of these strategies can realize the ESGD. Here we only concentrate on the polymorphic setup, since later we will focus on the simplest genotype-phenotype mapping, namely, when only two alleles at an autosomal locus determine the phenotypes. Observe that in the polymorphic setup, the subpopulations are considered as “players” (cf. Garay and Varga, 2003), since Definition 1 says that when the composition of the population falls in a neighbourhood of the ESGD, then there is no subpopulation having higher average fitness than the subpopulation in ESGD.

In the evolution coexisting types must have the same fitness. In our setup this also holds true. In Appendix B, we show that the equilibrium condition (Nash equilibrium, in game theoretical terms) can be given in the following form.

#### Equilibrium condition

If *x*\* ∈ int *S*_3_ is an ESGD and *x* is an arbitrary state in *S*_3_ then

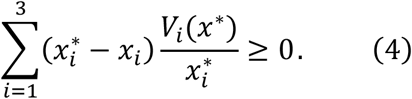

Moreover, one of the characteristic properties of interior Nash equilibria in matrix games, which gives a tool for finding interior Nash equilibria, remains true in this model, too. Namely, in interior Nash equilibria the fitness of different genotypes must be the same. Formally, let *x*\* ∈ int *S_m_* be an ESGD. Then, for every *i, j* ∈ {1, 2, 3} we have

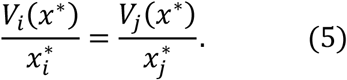

We note that equation (5) implies that an ESGD is a rest point of the genotype dynamics (6), see the next section.

### 2.3 Genotype dynamics

Now, let us follow the standard biological reasoning applied for the derivation of the replicator dynamics (see Garay et al., 2019, SI C). In the next generation the genotype distribution is

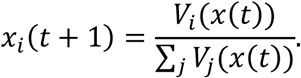

Therefore

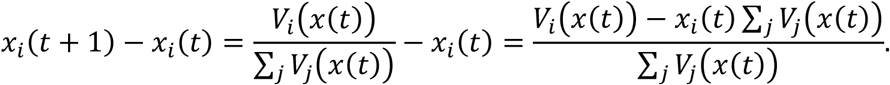

Hence, we derive the continuous-time model as follows:

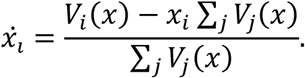

Clearly, 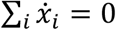 by ∑*_i_ x_i_* = 1. Since multiplying the right-hand side of the above equation by ∑*_j V_j__*(*x*) > 0 does not imply an essential change^1^, we get the following *genotype dynamics*:

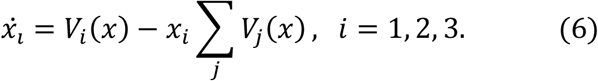

We emphasize that the genotype dynamics is quite different from the standard replicator dynamics, since according to the Mendelian inheritance, the offspring genotypes can be different from their parents’ genotypes. In contrast to the standard replication dynamics, the simplex *S*_3_ is only positively invariant under the above dynamics, in general. For instance, the homozygote vertices x_*_ = (1,0,0) and x_***_ = (0,0,1) are trivial rest points of the genotype dynamics, but x_**_ = (0,1,0), is not a rest point. We note that x_**_ is a rest point of the genotype dynamics (6) if and only if both homozygotes are lethal.

Observe that the dynamics in int *S*_3_ can be rewritten in the following “replicator dynamics” form:

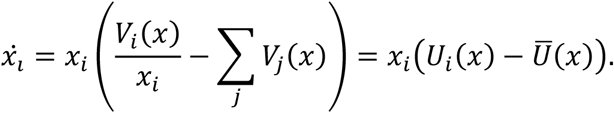

Applying the argumentation in Hofbauer and Sigmund (1998) we can prove the following result (see Appendix C for a general version).

#### Theorem 1.

*Assume that* x* ∈ int S_3_ *is an ESGD. Then x* is a locally asymptotically stable rest point of dynamics* (6).

## 3. Matrix game between siblings

Above we considered general models and did not fix where *n*_*k*(*ij*)_ came from. In the following application of our results we concentrate on matrix games. For simplicity, here we consider a two dimensional matrix game within each family, with a payoff matrix *A* ∈ *R*^2×2^. The phenotype distribution will be denoted by 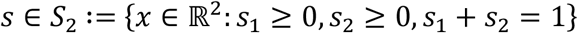. We assume that each individual follows a pure strategy, i.e., either *s*_1_ = (1,0) or *s*_2_ = (0,1). We emphasize that the entries of the payoff matrix are survival probabilities and the game is not iterated.

Under our biological assumptions specified in Section 2.1, for the sake of simplicity we only consider two alleles. Thus there are three genotypes with the genotype distribution *x* ∈ S_3_.

### Genotype-phenotype mapping

Mendelian dominant-recessive inheritance for two alleles at a single locus is a classification on genotypes, as follows: recessive genotype *G*_1_ = ([*a*],[*a*]) has phenotype *s*_1_ = (1,0), and genotypes *G*_2_ = ([*a*],[*A*]) and *G*_3_ = ([*a*],[*A*]) have phenotype *s*_2_ = (0,1). Clearly, on one hand, Mendelian dominant-recessive inheritance is symmetric in the sense that the heterozygotes have the same phenotype. On the other hand, from the perspective of the phenotypic composition of the families, the dominant-recessive inheritance is not symmetric, since the dominant phenotype appears in more families than the recessive one (see Table 2). To handle this asymmetry, in this paper the recessive phenotype corresponds to the first pure strategy. Consequently, in our population genetics model, the following two phenotypic payoff matrices,

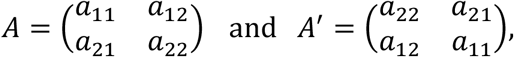

give different selection regimes, according to which pure strategy is the recessive one.

**Table 2.** Genotype survival table based on the mating table for matrix game, with phenotype *s*_1_ as recessive

| Genotypes of parents | Average number of couples | Average number of survived family members, for genotypes $G_1, G_2, G_3$ | | |
| --- | --- | --- | --- | --- |
| | | $G_1 \equiv ([a],[a])$<br>$s_1$ | $G_2 \equiv ([a],[A])$<br>$s_2$ | $G_3 \equiv ([A],[A])$,<br>$s_2$ |
| $G_1 \times G_1$ | $\frac{N}{2} x_1^2$ | $n_{1(11)} = n s_1 A s_1$ | 0 | 0 |
| $G_1 \times G_2$ | $N x_1 x_2$ | $n_{1(12)}$<br>$= \frac{n}{4} (s_1 A s_1 + s_1 A s_2)$ | $n_{2(12)}$<br>$= \frac{n}{4} (s_2 A s_1 + s_2 A s_2)$ | 0 |
| $G_1 \times G_3$ | $N x_1 x_3$ | 0 | $n_{2(13)} = n s_2 A s_2$ | 0 |
| $G_2 \times G_2$ | $\frac{N}{2} x_2^2$ | $n_{1(22)}$<br>$= \frac{n}{16} (s_1 A s_1 + 3 s_1 A s_2)$ | $n_{2(22)}$<br>$= \frac{n}{8} (s_2 A s_1 + 3 s_2 A s_2)$ | $n_{3(22)}$<br>$= \frac{n}{16} (s_2 A s_1 + 3 s_2 A s_2)$ |
| $G_2 \times G_3$ | $N x_2 x_3$ | 0 | $n_{2(23)} = \frac{n}{2} s_2 A s_2$ | $n_{3(23)} = \frac{n}{2} s_2 A s_2$ |
| $G_3 \times G_3$ | $\frac{N}{2} x_3^2$ | 0 | 0 | $n_{3(33)} = n s_2 A s_2$ |

Within each family the game theoretical interaction is well mixed. Consider an arbitrary type family and choose a child uniformly at random. Then the type of the chosen child and the type of their opponent in the matrix game are independent. This may seem surprising at first sight, since nobody can play with themselves. However, let us number the children from 1 to *n* in the order of birth time or in an arbitrary other way; the only requirement is that the ordering is independent of the genotype. Then we select the interacting pairs by a draw. Fix a pair, then the types of the players are clearly independent. For instance, in a *G*_2_×*G*_2_ family there are genotypes *G*_1_, *G*_2_ and *G*_3_, with probabilities ¼, ½ and ¼, and, according to the recessive-dominant inheritance, with phenotypes (pure strategies) *s*_1_ *s*_2_ and *s*_2_, respectively. Thus a *G*_1_ juvenile interacts with phenotypes *s*_1_ and *s_2_* with probabilities ¼ and ¾, so the survival rate of a focal *G*_1_ juvenile is 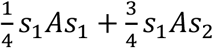. Moreover, the birth rate of a *G*_1_ juvenile in a *G*_2_×*G*_2_ family is ¼, thus the average number of surviving *G*_1_ juveniles in *G*_2_×*G*_2_ families is 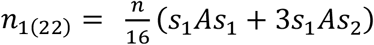. (For more details see Appendix D.)

Based on Table 2, the total number of individuals of genotypes *G*_1_, *G*_2_ and *G*_3_ in the next generation can be given as follows:

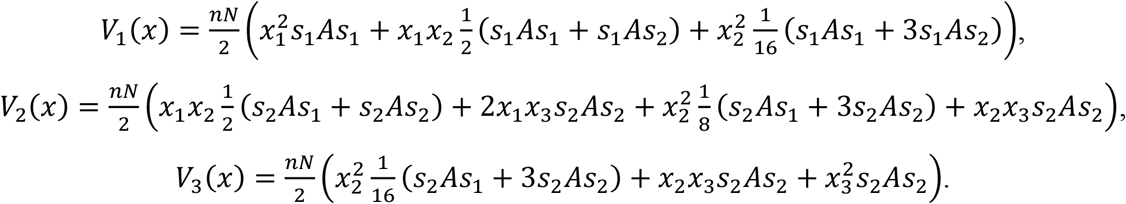

During our numerical investigations we focus on the genotype-phenotype mapping. In the next two examples we consider the same phenotypic selection situation, but in both examples we consider two subcases according to which phenotype is recessive.

#### Example 1

Let us consider the a numerical realization of the weak (cf. non-iterated) prisoner’s dilemma (i.e. *a*_21_ > *a*_11_ > *a*_22_ > *a*_12_, and 2*a*_11_ < *a*_21_ + *a*_12_), with the following phenotypic payoff matrices:

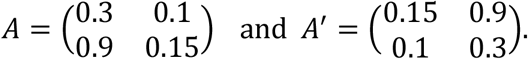

Observe that from the population genetics point of view, the difference between phenotypic payoff matrices *A* and *A’* is rooted in the genotype-phenotype mapping. The only difference is that which one of the pure strategies is recessive, which corresponds to swapping of the entries *a*_11_, *a*_22_, and *a*_12_, *a*_21_.

First, we consider matrix *A* and the corresponding interior Nash equilibrium, i.e. the solution of 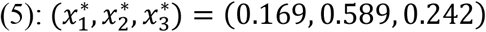. This interior Nash equilibrium *x*\* is evolutionarily stable, see Figure 1 that shows the function *F*(*x*, x*) around *x\**, where isoclines clearly indicate minimum.

**Figure 1.**
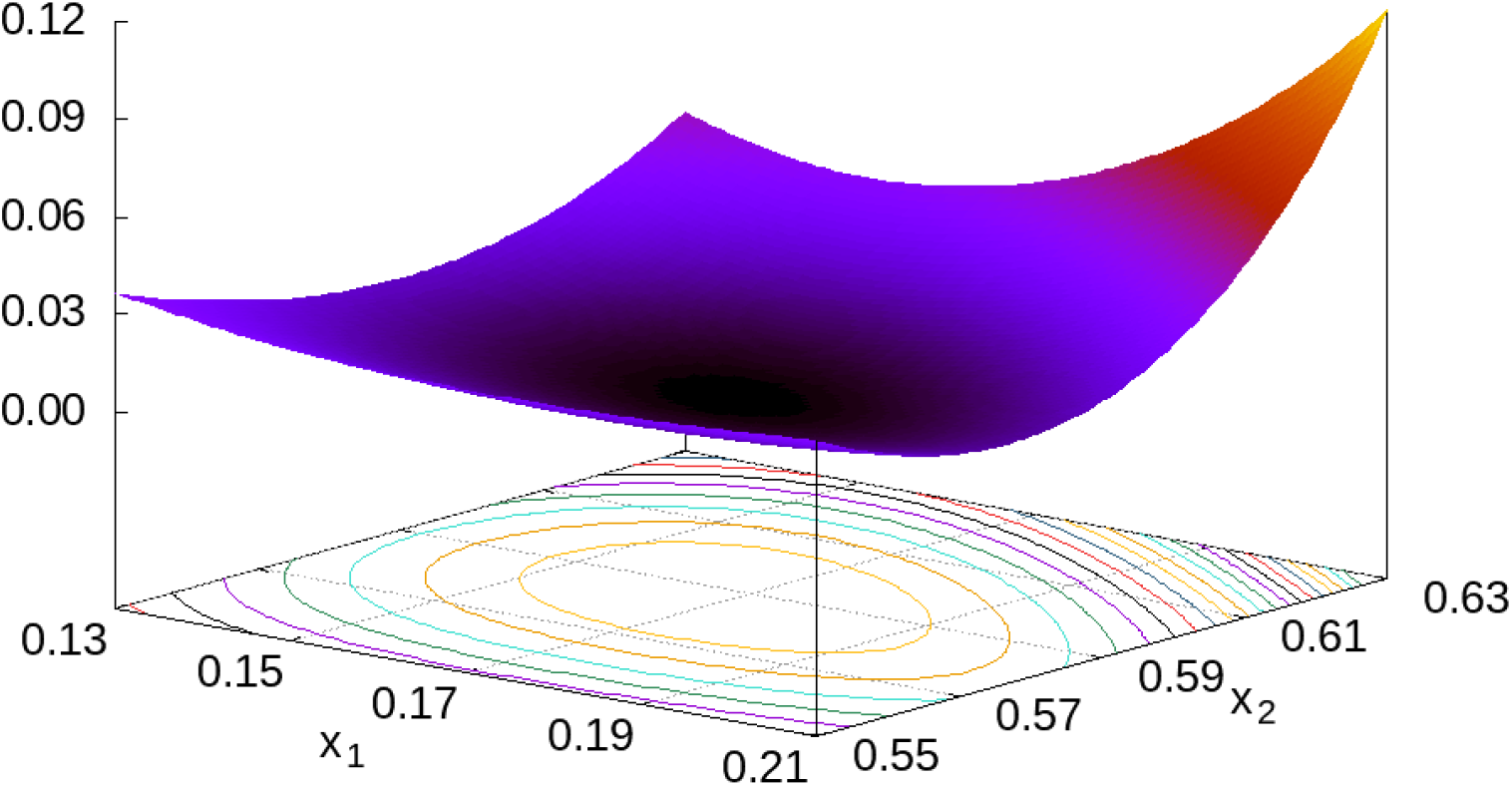
Function *F*(*x*, x*) around *x*\* = (0.169,0.589,0.242).

Second, for matrix *A*’ there also exists a mixed, but different ESGD, namely 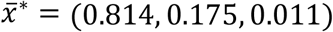. This shows that the coordinates of the mixed ESGD changes with the change of the genotype-phenotype mapping.

Now we present the visualization of Theorem 1. Figure 2 shows trajectories of genotype dynamics (6) on barycentric plots with matrix *A* (left panel) and with matrix *A*’ (right panel). In the first case, according to our numerical experiences, the interior rest point x* (the Nash equilibrium) is asymptotically stable, and the two trivial rest points (1,0,0) and (0,0,1) are unstable. By changing the genotype-phenotype mapping (the system is now governed by *A’*) the behavior of the two subcases do not change, i.e. there is a globally stable interior rest point in each case (since the function *F* has global minimum at *x*\*). However, the coordinates of these stable rest points are different.

**Figure 2.**
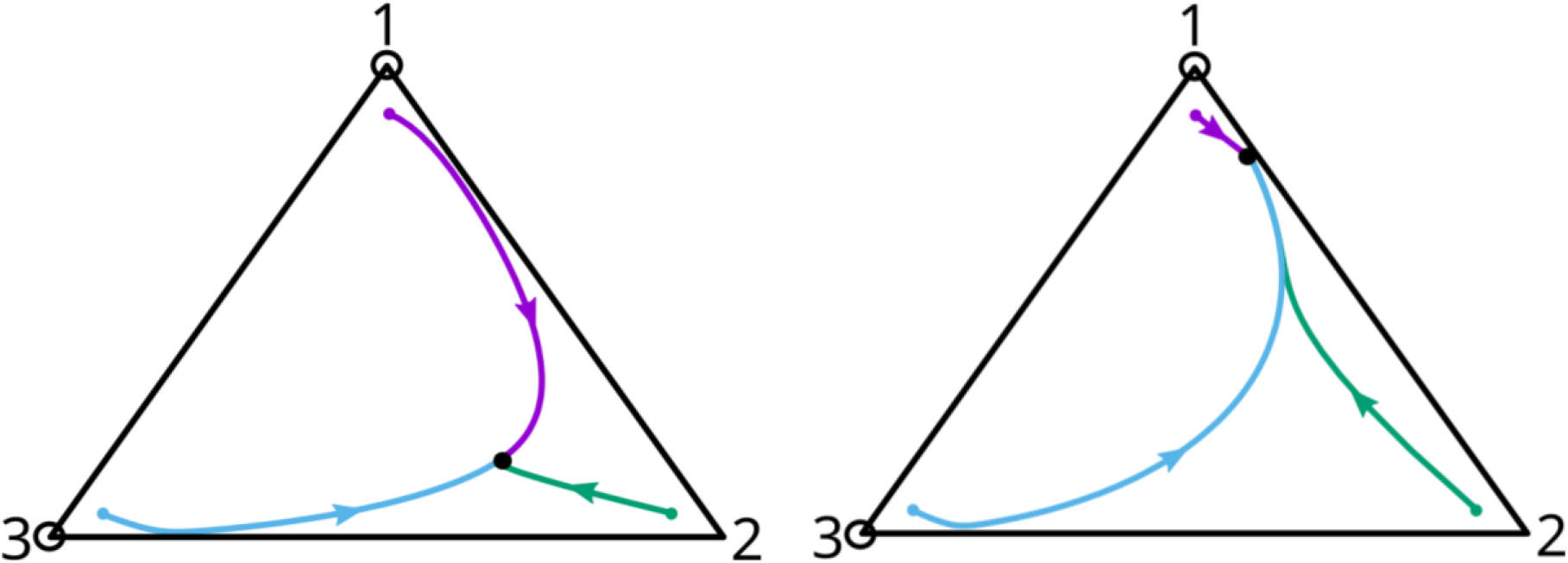
Trajectories of (6) on barycentric plot. **Left panel** The system is governed by phenotypic payoff matrix *A*. The globally asymptotically stable interior rest point is *x*\* = (0.169,0.589,0.242); *x*\*\* = (1,0,0) and *x*\*\*\* = (0,0,1) are unstable rest points. **Right panel** The system corresponds to the phenotypic payoff matrix *A’*. The new globally asymptotically stable interior rest point is 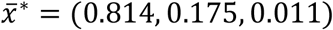; the unstable trivial rest points *x*\*\* and *x*\*\*\* are the same as for payoff matrix *A*. Black dots and circles denote asymptotically stable and unstable rest point, respectively. The initial conditions (denoted by small color dots) in both cases were (0.9, 0.05, 0.05), (0.05, 0.9, 0.05) and (0.05, 0.05, 0.9).

In summary, in the framework of familial selection model we have presented two numerical examples where the cooperator and defector phenotypes stably coexist; and the genotype-phenotype map can change the coordinates of the Nash equilibrium. We note that for the considered payoff matrices of the non-iterated prisoner’s dilemma, *A* and *A’*, there is only one and same pure classical ESS (as introduced by Maynard Smith and Price (1973) for asexual populations), namely, the defector pure strategy is a strict ESS. Thus the classical evolutionary game theoretical model and our population genetics model have quite different predictions.

#### Example 2

We also consider a phenotypic payoff matrix that corresponds to the prisoner’s dilemma game in the strong sense (cf. iterated prisoner’s dilemma, where *a*_21_ > *a*_11_ > *a*_22_ > *a*_12_ and 2*a*_11_ > *a*_21_ + *a*_12_, see matrix B).

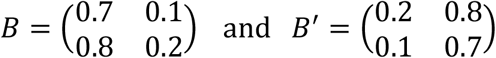

According to our numerical experiences, for matrix *B* there is no internal equilibrium, (1,0,0) is the global stable rest point of the system (pure cooperation). Changing the genotype-phenotype mapping by swapping the elements in the main diagonal (matrix *B’*) the global rest point becomes (0,0,1) (pure defector). In biological terms, we find that if the cooperator phenotype is recessive (matrix B) or dominant (matrix *B’*), then the cooperator and the defector phenotypes are the unique stable phenotypes, respectively. We note that the effect of genotype-phenotype mapping cannot be stronger than what we have found, in the sense that here the genotype-phenotype mapping determines which allele is eliminated by natural selection. The biological reason behind our observation is that the genotype-phenotype mapping determines the phenotypic composition of the families. When the altruistic allele is dominant, the altruistic phenotypes are more frequent in families than in the case where this allele is recessive. Thus the defector phenotype more often takes benefit from its cooperative siblings when the altruistic allele is dominant.

Finally, we also note the following. As it is well-known in the biological folklore, group selection theory has at least two ingredients, namely, a) the synergetic effect of the number of cooperators in the group on the individual fitness of the group members, b) the formation process of the groups. In our population genetic model, without the synergetic effect of group selection, the cooperator phenotype can exist. It seems that the group formation process is the really important factor in the evolution of cooperation.

From the Examples above the following two questions arise.

#### Question 1

What is the stability condition for the cooperator homozygote, according that the co-operator phenotype is recessive or dominant, when a matrix game gives the phenotypic selection? The detailed mathematical answer is in Appendix D1. In the main text we only mention the non-degenerated cases. The recessive cooperator homozygote is ESGD if

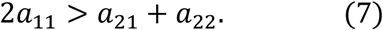

The dominant cooperator homozygote is ESGD if

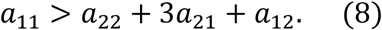

#### Question 2

What is a necessary mathematical condition for the coexistence of the two phenotypes? Clearly, the homozygote states are the only possible rest points of genotype dynamics (6) on the border of the simplex *S*_3_. Thus, according to the Poincaré–Bendixson theorem, if these two homozygote rest points are repellors, then (in biological terms) we have either stable or cyclic coexistence of the two phenotypes. The recessive cooperator homozygote is a repellor (see Appendix D1, Theorem 4), if

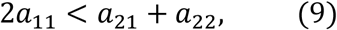

and the dominant cooperator homozygote is a repellor (see Appendix D1, Theorem 5), if

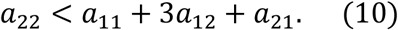

Thus inequalities (9–10) are sufficient for coexistence (see Appendix D1 for more details).

Finally, we have two remarks. 1. Our results are not in harmony with the standard view of inclusive fitness (e.g. Levin and Grafen, 2021). Firstly, the condition of stability of cooperator strategist varies according that the cooperator allele is recessive or dominant (see Examples 1–2). Secondly, the genetic relatedness (in monogamous population all individuals are full sibs) does not always occur in our conditions, see inequalities (8) and (10). 2. The standard group selection models focus on asexual populations (e.g. Nowak et al., 2010), thus these models cannot handle the effect of the genotype-phenotype mapping.

## 4. Discussion

Our model strictly belongs to the classical population genetics models, since the phenotypes of the individuals are exclusively determined by their genotypes. We concerned ourselves with the problem that in a given monogamous family what kind of phenotype can survive provided that the survival rate of each sib is determined by a phenotypic game between full siblings. The details of the genetic system determine the genotypic composition of each family (Garay et al., 2023), and the genotype-phenotype map yields the phenotypic composition. Our main observation is that the mathematical methods developed in classical, asexual matrix game theory (e.g. Hofbauer and Sigmund, 1998) can be useful in the population genetics models; namely, the static definition of ESGD implies the local stability of the genotype dynamics (6).

During the applications, we consider the simplest genetic system: the recessive-dominant Mendelian inheritance with one autosomal locus with two alleles and the prisoner’s dilemma takes place between siblings. By numerical studies, we found examples for the coexistence of the cooperator and the defector phenotypes. Our population genetics model calls again the attention to that the genotype-phenotype mapping has an important effect in familiar selection (Garay et al., 2019; Toro et al., 1982; Uyenoyama and Feldman, 1980), not only in the case of pure ESGD but also when there is a mixed ESGD. We note that, according to our knowledge, neither the kin selection nor the multilevel selection models have pointed out that in the evolutionary success of cooperative strategy the genotype-phenotype mapping has a crucial role, but our population genetic model show this.

From the perspective of kin and group selection theory, our monogamous population genetics model on familial selection can be considered as a special case study, because in our investigations only full siblings can have influence on the survival of others, since the interaction happens genetically well-defined groups, i.e., within families. However, our model is one of the simplest biological models for kin selection in diploid population, thus any general kin selection theory must contain our model as a special case. Moreover, in the overwhelming majority of eusocial animals the family defines the group (see e.g. naked mole-rat, honey bees), moreover in human societies the family is the basic, traditional social unit (Geary and Flinn, 2001). Based on that, we think the familial selection is an important issue to understand the evolutionary root of human altruism and cooperation, too. Furthermore, our population genetics model can handle the basic ideas of the kin and the group selection theories at the same time, i.e. the genetic relatedness and the group effect on survival probability of sibling together determine the evolutionary stability in genotype centred population genetics model of familiar selection (Garay et al., 2019, 2023). The speciality of familial selection is that the interactions only take place within the families, thus an open question arises: What will be the endpoint of such a selection regime when the game theoretical conflicts take place simultaneously within the family and between the members of different families, which can either belong to the same group or to different groups, too? On this research line the multilevel selection model might be incorporated into the population genetics theory in the future.

## Author contribution

**Conceptualization**: József Garay

**Methodology**: József Garay, Tamás F. Móri,

**Formal analysis and investigation**: Villő Csiszár, Tamás F. Móri, Tamás Varga,

**Visualization**: András Szilágyi

**Writing - original draft preparation**: József Garay, András Szilágyi, Tamás Varga, Villő Csiszár, Tamás F. Móri

**Writing - review and editing**: József Garay, András Szilágyi, Tamás Varga, Villő Csiszár, Tamás F. Móri

**Supervision**: József Garay

## Appendix A. General selection situation

First, we need some new notations. Consider *m* genotypes, thus the distribution of genotype belongs to the *m*-dimensional simplex

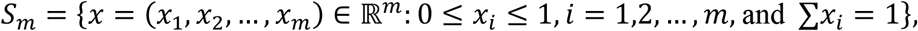

Let the probability of mating of genotypes *i* and *j* be *h_ij_* ∈ [0, 1]. In the applications of this paper we only consider panmixia, i.e. *h_ij_*(*x*) = *x_i_x_j_*.

Denote by *p*_*k*(*ij*)_ ∈ [0, 1] the probability that a pair of genotypes *i* and *j* has an offspring of genotype *k*. Clearly, the probability *p*_*k*(_ij_)_ can be calculated in the case of multi loci located at same or different chromosomes (with or without recombination). Obviously, in multi loci case the haploid stage is given by gametes. In the applications of this paper we only consider Mendelian inheritance with one locus and two alleles.

The phenotypic selection situation may be even more complicated. For instance, the offspring number may also depend on the genotypes of the parents, then let 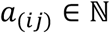 denote the number of offspring of a pair of genotypes *i* and *j*. We note the fecundity of a monogamous pair, *a*_(*ij*)_, can be determined by a frequency dependent game theoretical conflict between adults, too. Here we assume that *a*_(*ij*)_ = *n*. Moreover, let *q*_*k*(*ij*)_ ∈ [0, 1] be the probability of a survival of an offspring with genotype *k* and with parents of genotypes *i* and *j*. We note that, in applications, we concentrate on the additive model only, i.e., where a matrix game determines all *q*_*k*(*ij*)_. However, one can also consider non-additive models of the survival rate of siblings.

Now the total numbers of individuals of genotype *k* reads as

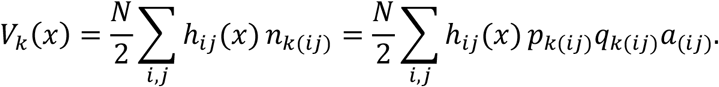

If *x* ∈ int *S_m_*, the average growth rate (fitness) of genotype *G_i_* is *U_i_*(*x*) = *V_i_*(*x*)/*x_i_*, and the average growth rate of the diploid population is 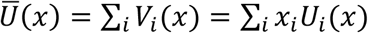. In order to make *Ū* strictly positive on the standard *m*-dimensional simplex we assume that *n*_*i*(*jk*)_ > 0 for all *i,j,k* ∈ {1, 2,…, *m*}. This in fact implies that *V_i_*(*x*) > 0 and *U_i_*(*x*) ≥ 0 for all *x* ∈ *s_m_* and *i* = 1, 2,…, *m*. In particular, *Ū*(*x*) > 0 for all *x* ∈ *S_m_*.

## Appendix B. Results on evolutionarily stable genotype distributions (ESGD)

We remind of the definition of evolutionary stability of a genotype distribution (see Definition 1). A genotype distribution *x*\* ∈ int *S_m_* is evolutionarily stable if

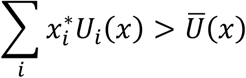

provided that x is sufficiently close to *x*\*. Consequently, *x*\* ∈ *S_m_* int is an ESGD if and only if

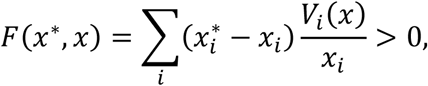

whenever *x* is close enough to *x*\*; that is, there is a *δ* > 0 such that the above inequality holds provided 0 < ║*x*\* – *x*║ < *δ*. Considering the Taylor expansion of *V_i_*(*x*)/*x_i_* at *x*\*, we have

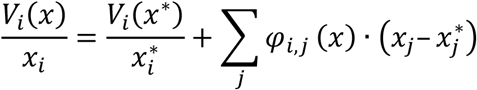

where *φ_i,j_*(*x*) = *∂_x_j__*(*V_i_*(*x*)/*x_i_*) + *ω_i,j_*(*x*) with 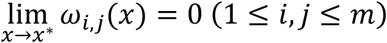, and there exist *K* and 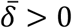, independent of *i* and *j*, such that |*φ_ij_*(*x*)| < *K* (1 ≤ *i,j* ≤ *m*) for any *x* with 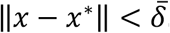. Since *S_m_* is bounded, there is a *t*_0_ such that 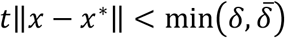 for every *x* ∈ *S_m_* and *t* ∈ (0, *t*_0_]. Consequently, if *x* ∈ *S_m_* and *t* ∈ (0, *t*_0_] then

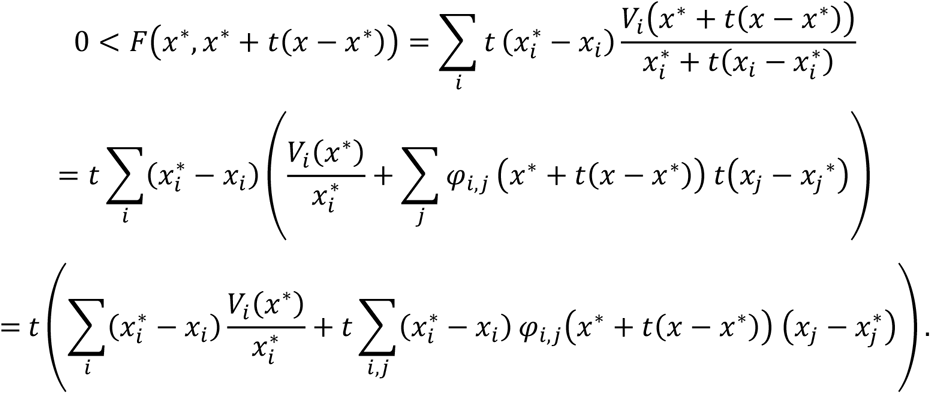

Dividing by *t* (recall that *t* > 0) we get that

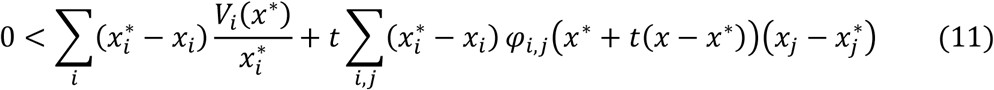

for any *t* ∈ (0, *t*_0_]. Since the functions *φ_i,j_* are uniformly bounded on the disk centered at *x*\* with radius 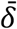, and the strict inequality above holds for arbitrary small positive *t*, we infer the following Nash equilibrium condition.

### Equilibrium condition

If *x*\* ∈ *S_m_* int is an ESGD and *x* is an arbitrary state in *S_m_* then

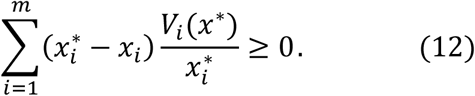

Moreover, one of the characteristic properties of interior Nash equilibria in matrix games is true in this model too, which gives a tool for finding interior Nash equilibria. This is the following.

#### Lemma 1.

Let *x*\* ∈ *S_m_* int be an ESGD. Then, for every *i,j* ∈ {1, 2,…, *m*} we have

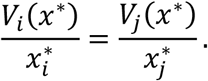

*Proof*. The proof follows the usual game theoretical reasoning (cf. (6.14)–(6.15) on p.64 in Hofbauer and Sigmund (1998)). Indeed, denote by *e_i_* the state *x* with *x_i_* = 1 and *x_j_* = 0 for *j* ≠ *i*, that is, the state in which every individual belongs to genotype *i*. According to the equilibrium condition (12) we have that

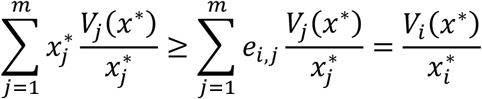

where *e_i,j_* is the *j*-th coordinate of *e_i_*, that is, *e_i,j_* = 1 if *j* = *i* and *e_i,j_* = 0 if *j* ≠ *i*. Multiplying by 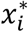 then summing from *i* = 1 to *i* = *m* we get

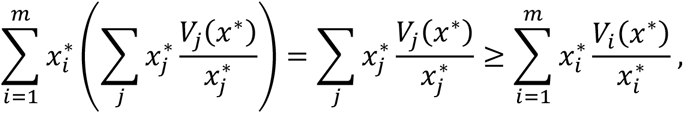

which is possible only if 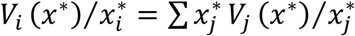 for every *i* = 1, 2,…, *m*. We have made use of the fact that 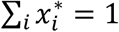.

We note that Lemma 1 implies that an ESGD is a rest point of the dynamics (6), see the next section.

Another consequence of the lemma is a negative semidefiniteness condition (cf. Haigh, 1975). By inequality (1), if *x* is close enough to *x*\*, then

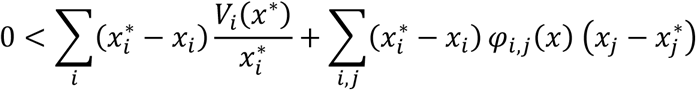

holds. On the other hand, we have 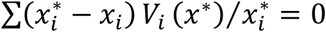 by Lemma 1, therefore this inequality simplifies to

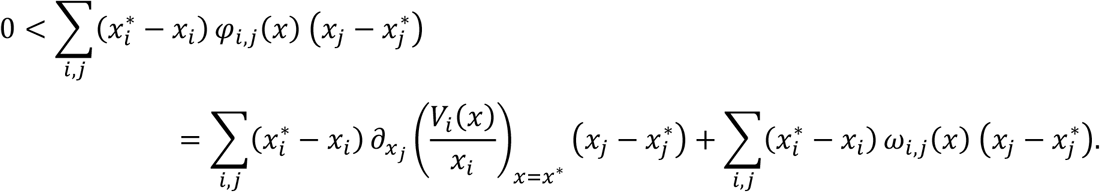

Since *ω_i,j_*(*x*) → 0 as *x* → *x*\* we infer that or

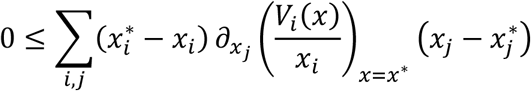

or

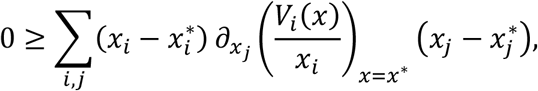

which just means that the matrix 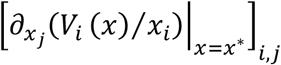 is negative semidefinite. (In the case of classical evolutionary matrix games negative definiteness holds because there the functions *ω_i,j_* are identically zero, see Exercise 6.4.3 in Hofbauer and Sigmund (1998)).

## Appendix C. Stability of ESGD with respect to genotype dynamics

Consider the genotype dynamics (6) derived in Section 2.3:

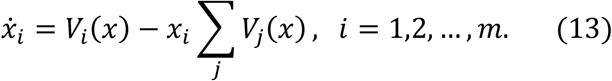

Here we prove the general version of Theorem 1.

### Theorem 1’.

*Assume that x*\* ∈ int *S_m_ is an ESGD. Then x* is a locally asymptotically stable rest point of dynamics* (13).

*Proof*. The proof follows the proof of Theorem 7.2.4 in Hofbauer and Sigmund (1998).

By Lemma 1, *x*\* is an interior rest point of the genotype dynamics (13).

We prove that *x*\* asymptotically stable. To see this, consider the function

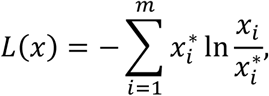

which is a Lyapunov function with respect to genotype dynamics (3) at state *x*\*. Here *L*(*x*\*) = 0 and *x*\* is a strict local minimum point of *L*, see the proof of Theorem 7.2.4 in Hofbauer and Sigmund (1998). On the other hand, the derivative of *L*(*x*) along the differential equation (13) is

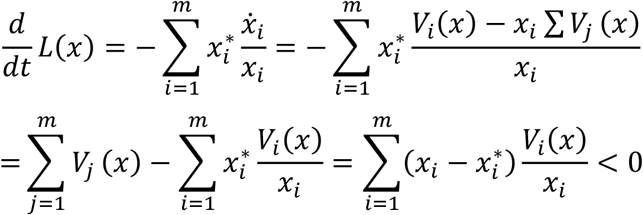

since *x*\* is an ESGD. This means that 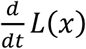 is negative definite at *x*\*. By Lyapunov’s theorem on stability, this implies that *x*\* is an asymptotically stable rest point of the genotype dynamics (13), see e.g. Theorem 2.6.1 in Hofbauer and Sigmund (1998); p.194 in Hirsch et al., (2004) or Theorem 3.5.1(b) in Kong (2014).

## Appendix D. Matrix game within a monogamous family

There are *N* individuals (*N* is a very large even integer), half of them are male, half of them are female. The proportions of the three genotypes *G*_1_ = [*a, a*], *G*_2_ = [*a, A*], *G*_3_ = [*A, A*] are the same among males and females, namely *x*_1_, *x*_2_, *x*_3_. They form *N*/2 male-female pairs (families) uniformly at random. Each family produces *n* offspring according to the Mendelian rules (*n* is supposed to be constant and even). Then these offspring form *n*/2 pairs uniformly at random, and the pairs play a matrix game with payoff matrix *A* ∈ ℝ^2×2^. If a player uses pure strategy *v* against an opponent using pure strategy *w*, then its survival probability will be *a_v,w_*(*v, w* ∈ {1, 2}). Of course, mixed strategies are also allowed. Suppose genotype *G*_1_ uses strategy vector *s*_1_ while the other genotypes use *s*_2_. Let *n*_*k*(*i,j*)_ denote the expected number of surviving offspring in a family of type *G_i_* × *G_j_*(*i, j, k* ∈ {1, 2, 3}). Firstly, we are going to compute these numbers.

During the calculus the following elementary facts will be used.

### Fact

Suppose that in a random experiment there are *m* possible mutually exclusive outcomes with corresponding probabilities by *π*_1_,…, *π_m_*, respectively. Let *ξ_i_*, indicate the number of times outcome number *i* is observed over *n* independent trials. Then the joint distribution of the random vector (*ξ*_1_,…, *ξ_m_*) is multinomial with parameters *n, π*_1_,…, *π*_m_. Moreover,

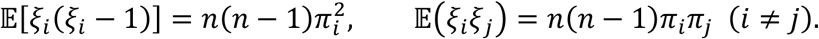

### Family type *G*_1_ × *G*_1_

Then *m* = 1, and *n*_1(11)_ = *ns*_1_*As*_1_, *n*_2(11)_ = *n*_3(11)_ = 0.

### Family type *G*_1_ × *G*_2_

Then *m* = 2, *π*_1_ = *π*_2_ = ½, and

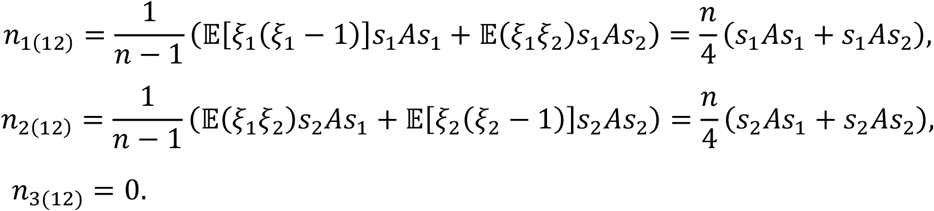

### Family type *G*_1_ × *G*_3_

Then *m* = 1, and *n*_2(13)_ = *nS*_2_*As*_2_, *n*_1(13)_ = *n*_3(13)_ = 0.

### Family type *G*_2_ × *G*_2_

Then *m* = 3, *π*_1_ = *π*_3_ = ¼, *π*_2_ = ν, and

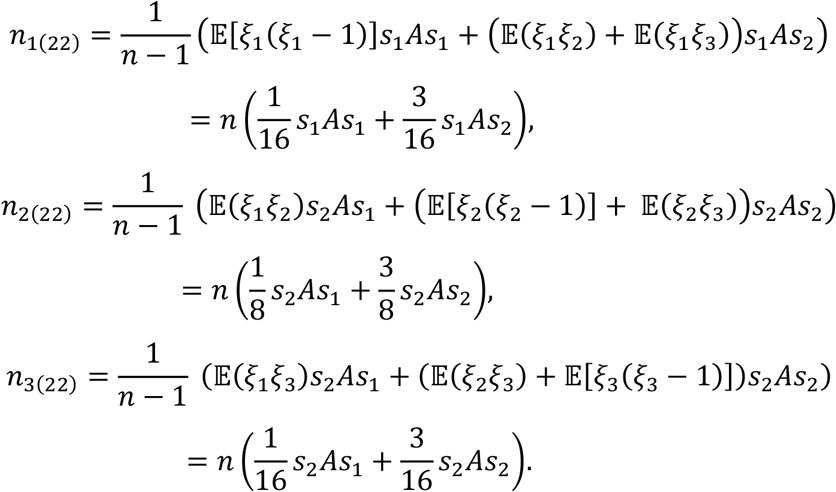

### Family type *G*_2_ × *G*_3_

In this case, the payoff is *s*_2_*As*_2_ in every game. Hence

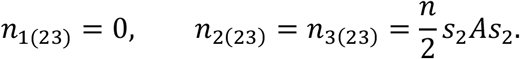

### Family type *G*_3_ × *G*_3_

Clearly, *n*_3(33)_ = *nS*_2_*As*_2_, *n*_1(33)_ = *n*_2(33)_ = 0.

Another, simpler way of computing these quantities can be found upon the following observation. Consider an arbitrary type family and choose a child uniformly at random. Then the type of the chosen child and the type of their opponent in the matrix game are independent. This may seem surprising at first sight, since nobody can play with themselves. However, let the children be numbered from 1 to *n* in the order of birth time or in an arbitrary other way; the only requirement is that the ordering is independent of the genotype. Then select the interacting pairs by a draw. Fix a pair, then the types of the players are clearly independent.

## Appendix D1. When will genotypes *G*_1_ and *G*_3_ be evolutionarily stable?

First, we are looking for conditions guaranteeing that the proportion *x*_1_ of genotype *G*_1_ increases from generation to generation provided it was sufficiently close to 1 in the beginning.

Let us apply Theorem 1 of Garay et al. (2019) and the subsection Application thereafter. Since we now consider non-overlapping generations, the survival probabilities of parents (denoted by *q*_*k*(*ij*)_ in that paper) are set to zero. Therefore we get that *G*_1_ is evolutionarily stable if any of the following conditions is satisfied.

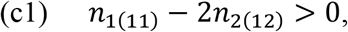

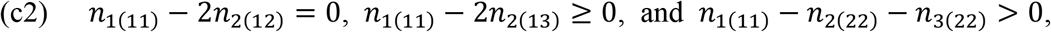

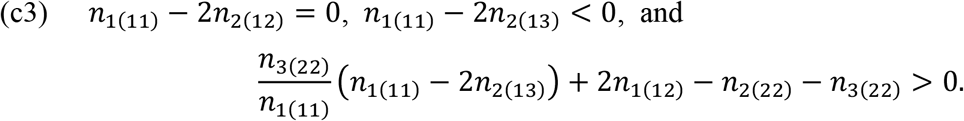

Let us plug in here the expressions we got for the quantities *n*_*k*(*ij*)_. For the sake of simplicity let us denote *s_i_As_j_* by *a_ij_*. Then

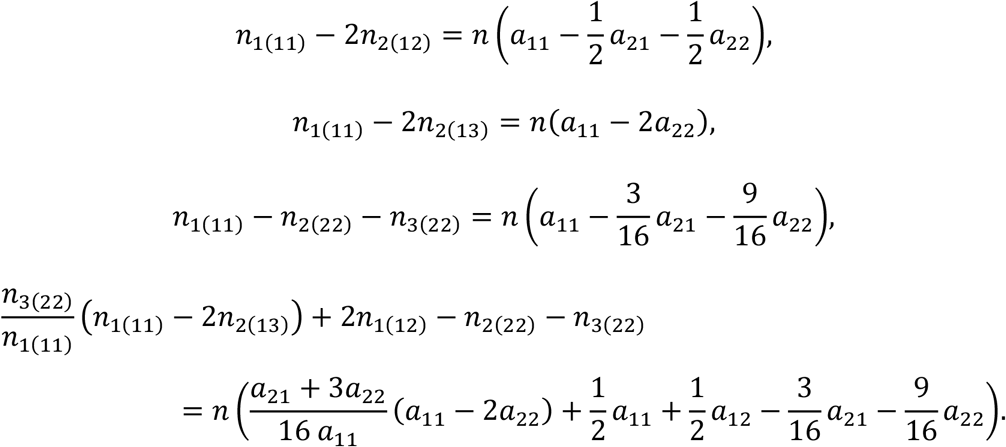

Thus, conditions (c1)–(c3) can be written in the following form.

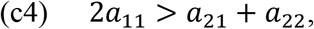

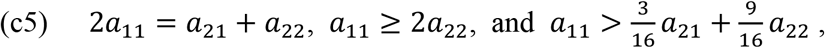

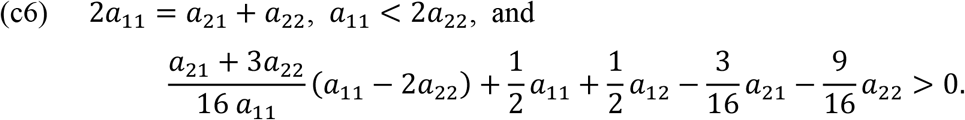

Replace *a*_21_ with 2*a*_11_ – *a*_22_ in the last inequality of (c5) to obtain 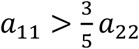. Since *a*_22_ > 0, this already follows from the second inequality. In (c6), after some calculus, the last inequality can be transformed into the more symmetric one (*a*_11_ + *a*_12_)(*a*_21_ + *a*_22_) > (*a*_11_ + *a*_22_)^2^. Consequently, we arrive at the following theorem.

### Theorem 2.

*Gis evolutionarily stable if the payoff matrix satisfies any of the following conditions.*

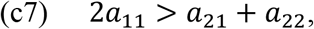

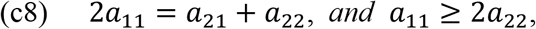

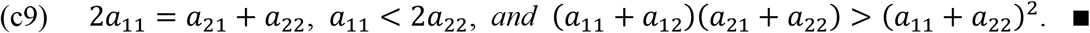

Let us turn to genotype *G*_3_. Conditions analogous with (c1)–(c3) can be obtained by interchanging the numbers 1 and 3 in the subscripts.

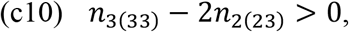

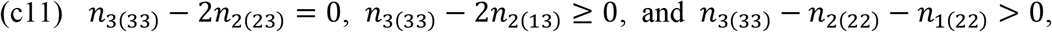

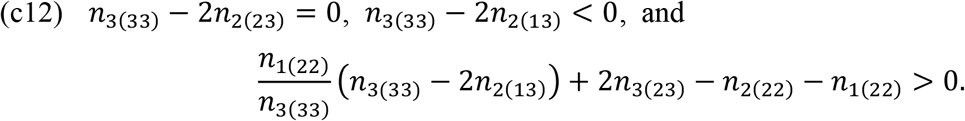

Here

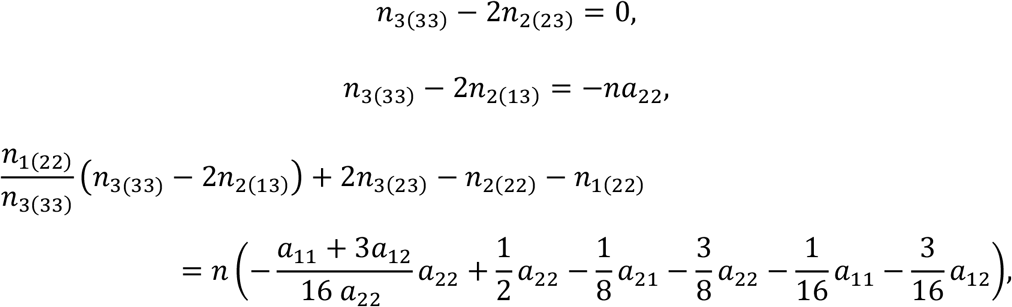

thus conditions (c10) and (c11) cannot hold, and condition (c12) reads *a*_22_ > *a*_11_ + 3*a*_12_ + *a*_21_.

### Theorem 3.

*G*_3_ *is evolutionarily stable if the payoff matrix satisfies *a*_22_ > *a*_11_ + 3*a*_12_ + *a*_21_*.

Evolutionary stability of *G*_1_ and *G*_3_ means that the vertices (1,0,0) and (0,0,1) are attractors, respectively. On the other hand, if in the conditions the last strict inequalities hold in the opposite direction, the corresponding vertex becomes a repellor (see Garay et al., 2019) That is, (1,0,0) is a repellor if any of the following conditions hold.

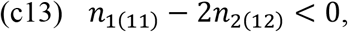

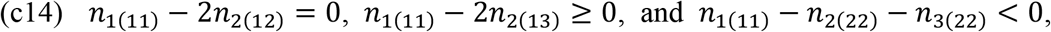

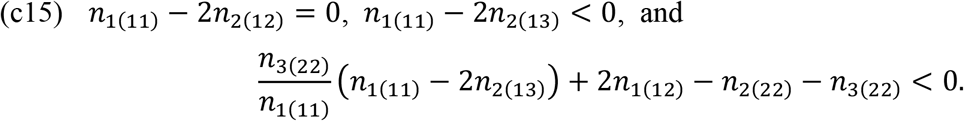

In terms of the payoff matrix we have

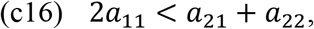

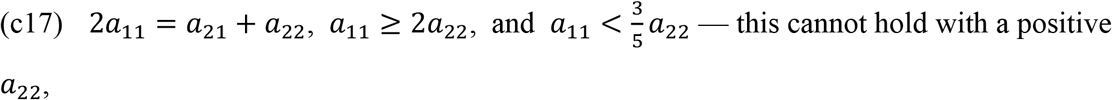

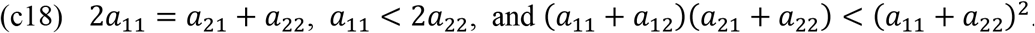

This latter will be satisfied if 3*a*_22_ ≥ *a*_11_ + 4*a*_12_, because then

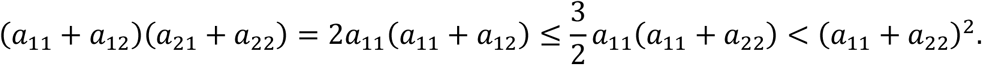

### Theorem 4.

(1,0,0) *is a repellor if either* 2*a*_11_ < *a*_21_ + *a*_22_, *or* 2*a*_11_ = *a*_21_ + *a*_22_, *a*_11_ < 2*a*_22_, *and* 3*a*_22_ ≥ *a*_11_ + 4*a*_12_.

Similarly, (0,0,1) is a repellor if any of the following conditions hold.

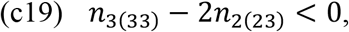

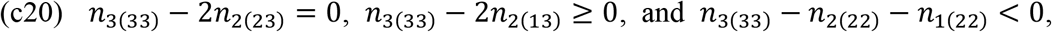

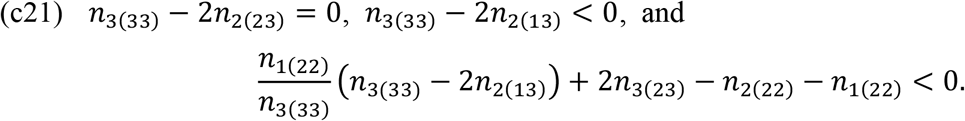

As we have already seen, (c19) and (c20) can never hold. In terms of the payoff matrix (c21) means that *a*_22_ < *a*_11_ + 3*a*_12_ + *a*_21_.

### Theorem 5.

(0,0,1) *is a repellor if a*_22_ < *a*_11_ + 3*a*_12_ + *a*_21_.

## Appendix D2. How many identical copy do a focal gene produce?

1. First, let the focal gene be [*a*]. Let us select a gene uniformly at random from all paternal [*a*] genes (by symmetry, we can confine ourselves to paternal genes). Then it is

1.1. in an individual of genotype *G*_1_ with probability 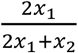. Then his spouse is

1.1.1. of genotype *G*_1_ too with probability *x*_1_. In this case all offspring are of the same genotype and half of the surviving offspring inherit the focal gene on the average: 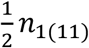.
1.1.2. of genotype *G*_2_ with probability *x*_2_. In this case again half of the surviving offspring inherit the focal gene on the average: 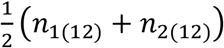.
1.1.3. of genotype *G*_3_ with probability *x*_3_. In this case again half of the surviving offspring inherit the focal gene on the average: 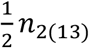.
1.2. in an individual of genotype *G*_2_ with probability 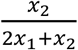. Then his spouse is

1.2.1. of genotype *G*_1_ with probability *x*_1_. In this case all type *G*_1_ offspring inherit the focal gene and only they do: *n*_1(12)_.
1.2.2. of genotype *G*_2_ with probability *x*_2_. Again, in this case all type *G*_1_ offspring inherit the focal gene, but there can be type *G*_2_ (and even type *G*_3_) offspring too. The latter ones do not inherit the focal gene, while about half of the the former ones do: 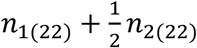.
1.2.3. of genotype *G*_3_ with probability *x*_3_. In this case all type *G*_2_ offspring inherit the focal gene and only they do: *n*_2(23)_. Thus the average number of identical copies of the focal [α] gene in the progeny is equal to

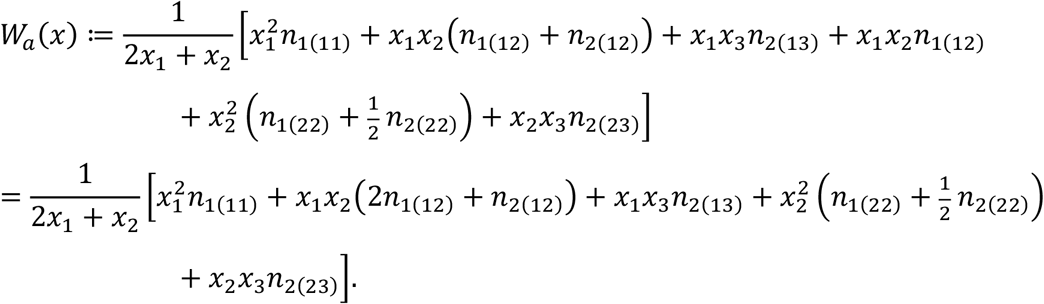
2. The average number of identical copies of a focal [Λ] gene in the progeny can be computed in a similar way. The focal gene is

2.1. in an individual of genotype *G*_2_ with probability 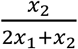. Then his spouse is

2.1.1. of genotype *G*_1_ with probability *x*_1_. In this case all type *G*_2_ offspring inherit the focal gene and only they do: *n*_2(12)_.
2.1.2. of genotype *G*_2_ with probability *x*_2_. In this case all type *G*_3_ offspring inherit the focal gene, but there can be type *G*_2_ (and even type *G*_1_) offspring too. The latter ones do not inherit the focal gene, while about half of the former ones do: 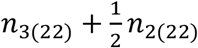.
2.1.3. of genotype *G*_3_ with probability *x*_3_. In this case all type *G*_3_ offspring inherit the focal gene and only they do: n_3(23)_
2.2. in an individual of genotype *G*_3_ with probability 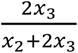. Then his spouse is

2.2.1. of genotype *G*_1_ with probability *x*_1_. In this case half of the surviving offspring inherit the focal gene on the average: 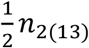.
2.2.2. of genotype *G*_2_ with probability *x*_2_. In this case again half of the surviving offspring inherit the focal gene on the average: 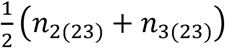.
2.2.3. of genotype *G*_3_ too with probability *x*_3_. In this case all offspring are of the same genotype and half of the surviving offspring inherit the focal gene on the average: 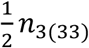.

Thus the average number of identical copies of the focal [Λ] gene in the progeny is equal to

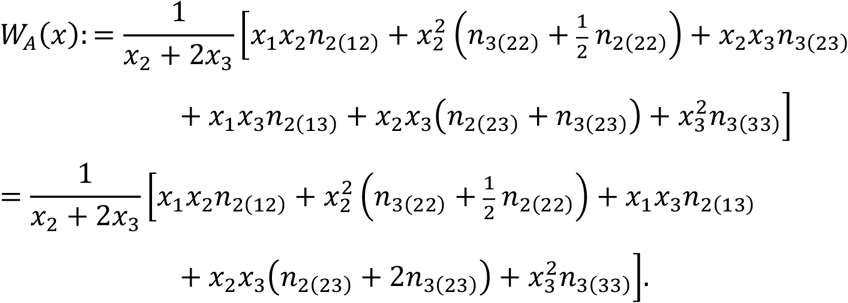

Note that these results can be obtained in another way. Denoting by *V_i_*(*x*) the number of individuals of genotype *G_i_*, in the next generation we clearly have

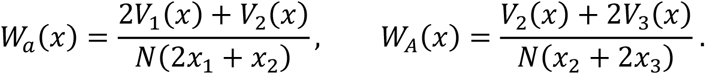

The denominator contains the total number of genes in the present generation that are of the same type as the focal one, while in the numerator there is the same quantity with respect to the next generation. This is the average number of identical copies of a focal gene in the progeny, indeed.

Obviously,

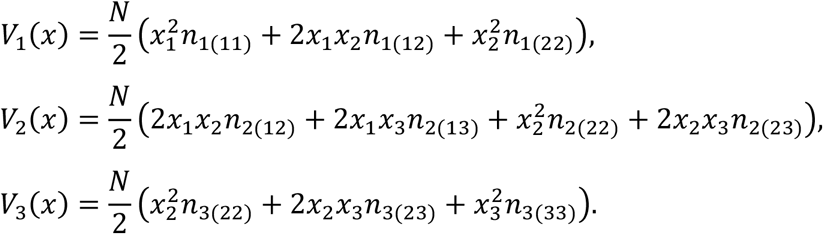

### Theorem 6.

*Suppose* 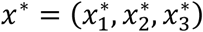 *is an ESGD. Then W_a_*(*x\**) = *W_A_*(*x\**).

*Proof*. By Lemma 1 we have

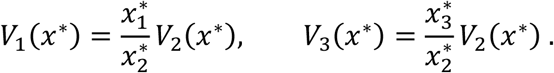

Therefore

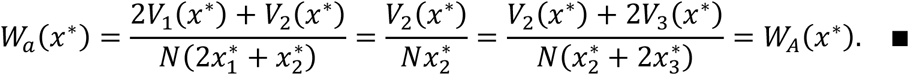

## Footnotes

1 This operation does not affect the orbits, only changes the velocity (see e.g. Hofbauer and Sigmund, 1998, pp. 118-119).

